# Effects of adaptive harvesting on fishing down processes and resilience changes in predator-prey and tritrophic systems

**DOI:** 10.1101/290460

**Authors:** Eric Tromeur, Nicolas Loeuille

## Abstract

Many world fisheries display a declining mean trophic level of catches. This “fishing down the food web” is often attributed to reduced densities of high-trophic-level species. We show here that the fishing down pattern can actually emerge from the adaptive harvesting of two- and three-species food webs, where changes in fishing patterns are driven by the relative profitabilities of the harvested species. Shifting fishing patterns from a focus on higher trophic levels to a focus on lower trophic levels can yield abrupt changes in the system, strongly impacting species densities. In predator-prey systems, such regime shifts occur when the predator species is highly valuable relative to the prey, and when the top-down control on the prey is strong. Moreover, we find that when the two species are jointly harvested, high adaptation speeds can reduce the resilience of fisheries. Our results therefore suggest that flexibility in harvesting strategies will not necessarily benefit fisheries but may actually harm their sustainability.

## Introduction

Many world fisheries have experienced over the 20th century a general pattern of reduced mean trophic level of catches, called “fishing down the food web” (Pauly et al. 1998). This phenomenon may be linked to different situations. One possibility is that higher (more commercially valuable) trophic levels are targeted and that their collapse leads to the exploitation of lower trophic levels. However, reduction of mean trophic levels of catches may also be linked to the progressive inclusion of more low trophic level stocks in the fishery (Essington et al. 2006, Branch et al. 2010), a pattern known as “fishing up the food web”. Decrease in mean trophic levels can also be expected if stocks are fished proportionally to their availability (Branch et al. 2010). Therefore a variety of situations lead to such “fishing down the food web”. In the present paper, we contend that fishing down patterns can be explained by the adaptive behaviour of fishermen faced with declining abundances of higher trophic levels.

Empirical data suggest that fishermen often act as adaptive foragers in ecosystems, optimizing the rate of encounter with resources (Bertrand et al. 2007), and regularly switching the species they fish to maximize profits (Acheson 1988). Adaptive harvesting can also result from the adaptive management of fisheries (Holling 1978), where management authorities, based on population abundances, change quota or effort limits, thereby altering the distribution of fishing efforts (Walters 1986).

Similar to adaptive foraging in food webs, adaptive harvesting is likely to affect the resilience of fisheries. Resilience is generally understood as the ability of an ecosystem to absorb changes and still persist (Holling 1973). Adaptive foraging has notably been shown to stabilize food web dynamics by reallocating predation pressures on more abundant species, thereby creating a stabilizing negative feedback (Loeuille 2010*a*). As a result, high adaptation speeds improve the resilience of complex food webs, as shown in (Kondoh 2003).

In the present work, we assess the consequences of adaptively harvesting a trophic community. To this aim, we first use a model of an adaptively harvested predator-prey community. We first investigate the effect of increasing total fishing effort on the fishing pattern. We then examine the impact of adaptive harvesting on the resilience of the fishery, by focusing on three aspects of resilience: the propensity to undergo regime shits, the ability of returning to equilibrium following a pulse perturbation, and the speed of this return. Our results suggest that adaptive harvesting results in the progressive decline in the mean trophic level of catches. Therefore, fishing down the food web naturally emerges from the adaptive harvesting in our model. Interestingly, our results also suggest that such an adaptive behaviour may be harmful for the overall resilience of the system. Finally, we numerically show similar results in a tritrophic community.

## Model

### Two-species fishery

We use a simple Lotka-Volterra predator-prey model, where both species are harvested:

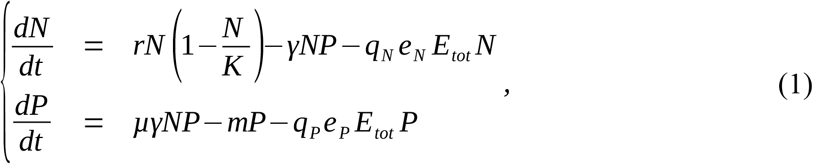

where *N* is prey density, *P* predator density, *K* the prey carrying capacity (all in ind/surface or volume), *r* the prey growth rate (per unit of time), *γ* the predator attack rate (per capita and per unit of time), *μ* the prey-to-predator conversion efficiency - that is the efficiency with which predators convert eaten prey into new predators (dimensionless), *m* the predator mortality (per unit of time), *q_N_* and *q_P_* the prey and predator catchabilities (per unit of time and of effort), *E_tot_* the total fishing effort, and *e_N_* and *e*_*P*_ the shares of effort allocated to prey and predator (dimensionless). These effort shares can represent the proportion of time spent using a prey- or predator-specific gear, or the proportion of vessels that specifically harvest prey or predators. Considering a single fishing vessel, the effort *E_tot_* may represent the number of hours spent at sea, while considering the entire fishery, the effort may represent the number of active fishing vessels. Based on the replicator equation, which is commonly used to model such evolutionary dynamics (Page and Nowak 2002) or to model adaptive foraging within food webs (Valdovinos et al. 2010), the share of effort allocated to prey *e_N_* varies following the equation:

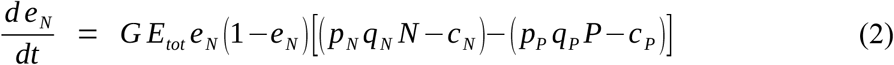

where *p*_*N*_ and *p_P_* are the prices of prey and predators, *c_N_* and *c_P_* are the per-unit-effort costs of harvesting prey and predators, and *G* is the adaptation speed. The share of effort allocated to predator is then *e*_*P*_ = 1−*e*_*N*_. Following Equation (2), the share of prey effort increases if the marginal utility of harvesting the prey (first term within brackets) is higher than the marginal utility of harvesting the predator (second term within brackets). The speed at which reallocation of the effort may occur (eg, technical constraints) is dependent on adaptation speed *G*. Note that equation (2) implicitly assumes that costs and prices are fixed (including independent of densities) when computing marginal utilities.

To assess whether our results depend on the linearity of the functional response in the initial system (1), and to allow population cycles to occur, we also investigate a predator-prey model with a Holling type-II functional response (Rosenzweig-MacArthur model):

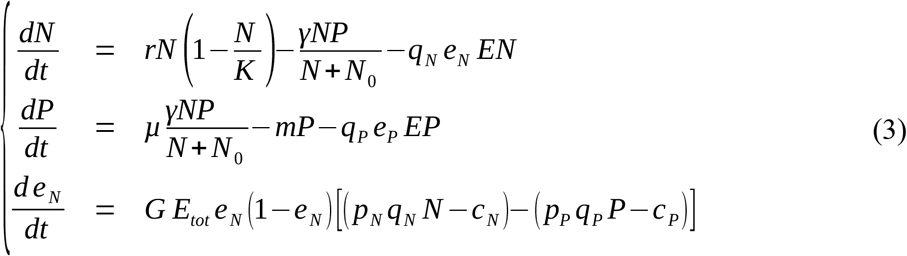

where *N*_0_ is the prey density at which the predator’s attack rate reaches half of its maximum.

### Local stability analysis and resilience calculation

The local stability of the system can be assessed by computing the eigenvalues of the Jacobian of the system at equilibrium. The general formulation of this Jacobian for the two-species model is:

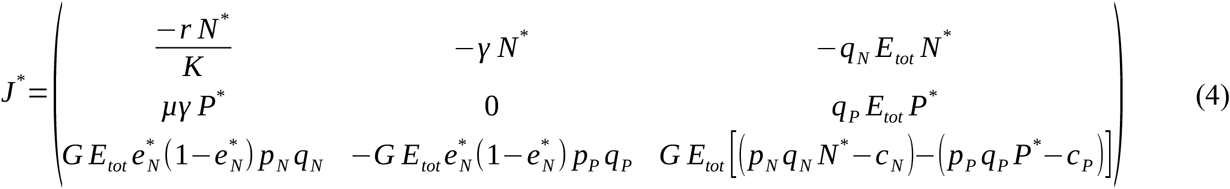

Resilience can be measured by the return time to equilibrium after a perturbation, small return times indicating high levels of resilience. To obtain this measure of resilience, we compute from the Jacobian matrix the eigenvalue with the largest real part *λ*_*m*_. The return time to equilibrium after a small perturbation is then τ=−1/ Re(λ_*m*_) (Loeuille 2010*b*).

### Three-species fishery

Finally, we tackle whether the result of the two species model (system (1)) is restricted to very simple, two species systems. We study adaptive harvesting in a three species food chains. We use a simple Lotka-Volterra tritrophic chain model, where all species can be harvested:

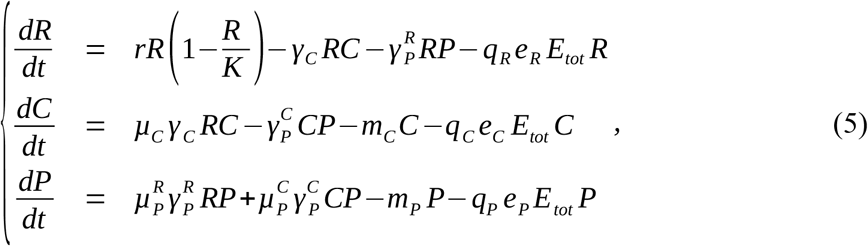

where *R* is resource density, *C* consumer density, *P* predator density, *r* the resource growth rate, *K* the resource carrying capacity, *γ*_*C*_ the consumer attack rate, 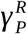 and 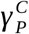 the attack rates of predators on resource and consumers, *μ*_*C*_, 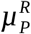 and 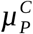 the resource-to-consumer, resource-to-predator and consumer-to-predator conversion efficiencies, *m*_*C*_ and *m_P_* the consumer and predator mortalities, *q*_*R*_, *q*_*C*_ and *q*_*P*_ the respective consumer, prey, and predator catchabilities, and *E_tot_* the total fishing effort. The shares of effort allocated to either of the three harvested species (*e*_*R*_, *e*_*C*_ or *e*_*P*_) vary following these equations:

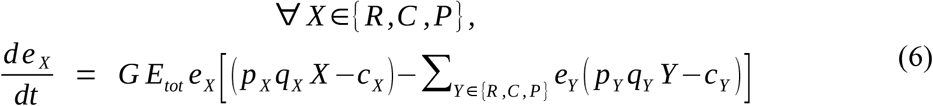

where *p*_*R*_, *p*_*C*_ and *p_P_* are the respective prices of resources, consumers and predators, and *c*_*R*_, *c_C_* and *c_P_* are the respective per-unit-effort costs of harvesting resources, consumers and predators. The relationship between the shares of total effort allocated to the three harvested species is then *e*_*R*_ + *e*_*C*_ +*e*_*P*_ =1. These effort shares can represent the proportion of time spent using a prey- or predator-specific gear, or the proportion of vessels that specifically harvest prey or predators. Following Equation (4), the share of effort allocated to a species increases if the marginal utility of only harvesting this species is higher than the marginal utility of harvesting the three species, weighted by their respective effort shares.

### Mean trophic level of catches

In the two-species model, the mean trophic level of catches (MTLC) is computed as follows :

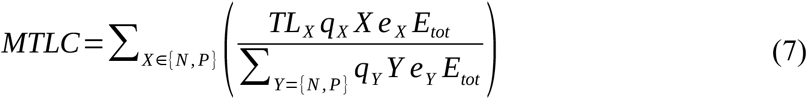

where the trophic levels of prey and predator are respectively *TL*_*N*_ = 2, and *TL*_*P*_=3. In the three-species model, the mean trophic level of catches (MTLC) is computed as follows :

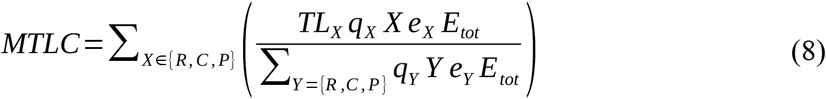

where the trophic levels of resource, consumer and predator are respectively *TL*_*R*_=1, *TL_C_*=2, and *TL*_*P*_=3.

## Results

### Two-species fishery

We first mathematically analyze the yield and ecological states of the system at equilibrium, corresponding to situations in which the time variations in system (1) and equation (2) are simultaneously null. We therefore compute equilibrium densities and effort shares so that *dN* / *dt* =0 *dP*/ *dt* =0 and *d e*_*N*_ / *dt*=0. Three equilibrium fishing patterns emerge: a predator-focused, a prey-focused and a mixed fishing pattern (see Figure A1 in Appendix A for illustration). All analytical expressions, as well as feasibility and stability conditions of these equilibria are shown in Table 1.

**Table 1.** Summary table of equilibria and their feasibility and stability conditions. Expressions of limit efforts 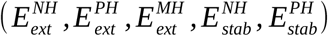 are found in the Appendix. 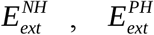 and 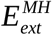 are the efforts above which the predator gets extinct in the prey-focused, predator-focused, and mixed harvest equilibria. 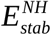 is the effort above which the prey-focused equilibrium is stable, while 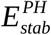 is the effort below which the predator-focused equilibria is stable. Only stability conditions that are not redundant with feasibility conditions are shown.

| Equilibrium | Predator harvest (PH) | Mixed harvest (MH) | Prey harvest (NH) |
| --- | --- | --- | --- |
| Analytical expressions | $N^* = \frac{m + q_P E_{tot}}{\mu \gamma}$ $P^* = \frac{r}{\gamma} \left( 1 - \frac{N^*}{K} \right)$ $e_N^* = 0$ | $N^* = \frac{m}{\mu \gamma}$ $P^* = \frac{1}{\gamma} \left[ r \left( 1 - \frac{m}{\mu \gamma K} \right) - q_N E_{tot} \right]$ $e_N^* = 1$ | $N^* = \frac{m + q_P (1 - e_N^*) E_{tot}}{\mu \gamma}$ $P^* = \frac{p_N q_N N^* + c_P - c_N}{p_P q_P}$ $e_N^* = \frac{1}{q_N E_{tot}} \left[ r \left( 1 - \frac{N^*}{K} \right) - \gamma P^* \right]$ |
| Feasibility conditions | $E_{tot} < E_{ext}^{PH}$ | if $\frac{q_P}{q_N} > \frac{\gamma K}{r} \left( \mu - \frac{p_N}{p_P} \right)$:<br>$E_{tot} < E_{ext}^{MH}$<br>$E_{stab}^{PH} < E_{tot} < E_{stab}^{NH}$<br>if $\frac{q_P}{q_N} < \frac{\gamma K}{r} \left( \mu - \frac{p_N}{p_P} \right)$:<br>$E_{tot} > E_{ext}^{MH}$<br>$E_{stab}^{NH} < E_{tot} < E_{stab}^{PH}$ | $E_{tot} < E_{ext}^{NH}$ |
| Stability conditions | $E_{tot} < E_{stab}^{PH}$ | $\frac{q_P}{q_N} > \frac{\gamma K}{r} \left( \mu - \frac{p_N}{p_P} \right)$ | $E_{tot} > E_{stab}^{NH}$ |

The predator-focused equilibrium can be written as follows:

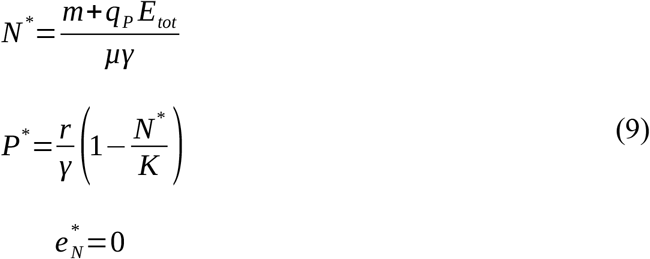

Stability analysis of this equilibrium is detailed in the Appendix. It shows that the equilibrium is stable provided *p_P_q_P_P*^*^−*c_P_* > *P_N_q*_*N*_*N*^*^−*c_N_*, which means that the marginal utility of harvesting the predator is higher than the marginal utility of harvesting the prey at equilibrium. Increasing fishing effort reduces predator density, and thereby enhances prey density. As a result, increasing fishing pressure improves the marginal utility of harvesting the prey, eventually destabilizing the predator-focused equilibrium (see Appendix A for detailed analytical calculations).

The prey-focused equilibrium can be written as follows:

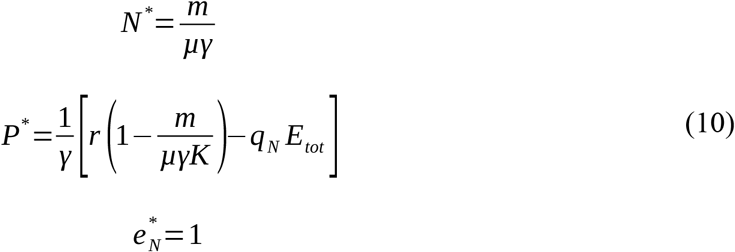

It is stable provided *p_P_q_P_P*^*^−*c_P_* > *P_N_q*_*N*_*N*^*^−*c_N_*, which means that the marginal utility of harvesting the prey is higher than the marginal utility of harvesting the predator. Increasing fishing effort on prey does not affect prey density, as prey is top-down controlled, ie its density depends on its predator demographic parameters. Increasing fishing effort however yields an indirect negative effect on predator density. Higher fishing pressure thus maintains the prey-focused fishing pattern, until the predator goes extinct (see Appendix A).

The mixed harvest equilibrium can be written as follows:

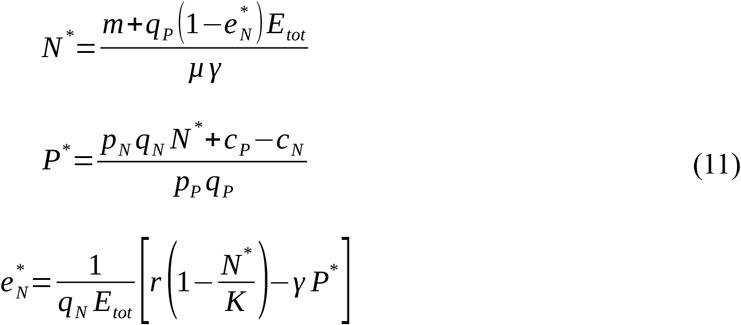

In this equilibrium the two species are simultaneously harvested because the marginal utility of harvesting the prey is equal to the marginal utility of harvesting the predator: *p_P_q_P_P*^*^−*c_P_* = *P_N_q*_*N*_*N*^*^−*c_N_*. This mixed fishing pattern is stable provided

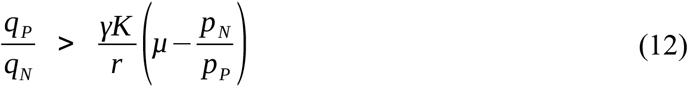

This condition is dependent on ecological, economic and technical parameters. It holds whenever the price of the prey is large enough relative to the price of the predator modulated by the conversion efficiency (*p*_*N*_ > *μ p_P_*). In particular, if the prey price is larger than the predator price, this stability condition always holds. Stability is also favored by a high predator-to-prey catchability ratio (*q*_*P*_ / *q*_*N*_), as well as by a low attack rate of the predator (*γ*), relative to the intraspecific competition rate of the prey (*r* / *K*). Therefore, the equilibrium is more likely to be stable when prey species are controlled by competition rather than by predation (i.e., in bottom-up controlled systems, a likely case in many fisheries).

The equilibrium fishing patterns depend on the the total fishing effort. As illustrated in Figure 1 (and analytically shown in Appendix A), the fishery shifts from a predator-focused harvest to a prey-focused harvest with increasing fishing efforts. In between, a stable mixed harvest of predator and prey is reached when condition (12) holds (Fig. 1a). In this case, the effort below which the predator-focused equilibrium is stable (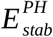, see definition in Appendix A) is lower than the effort above which the prey-focused equilibrium is stable (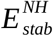, see definition in Appendix A). The progressive shift from a predator-focused to a prey-focused fishing pattern leads to a general reduction in predator density (Fig. 1c), and a decline in the mean trophic level of catches (Fig. 1e). As predator density decreases with increasing efforts to reach extinction, the mean trophic level of the community also decreases.

**Figure 1:**
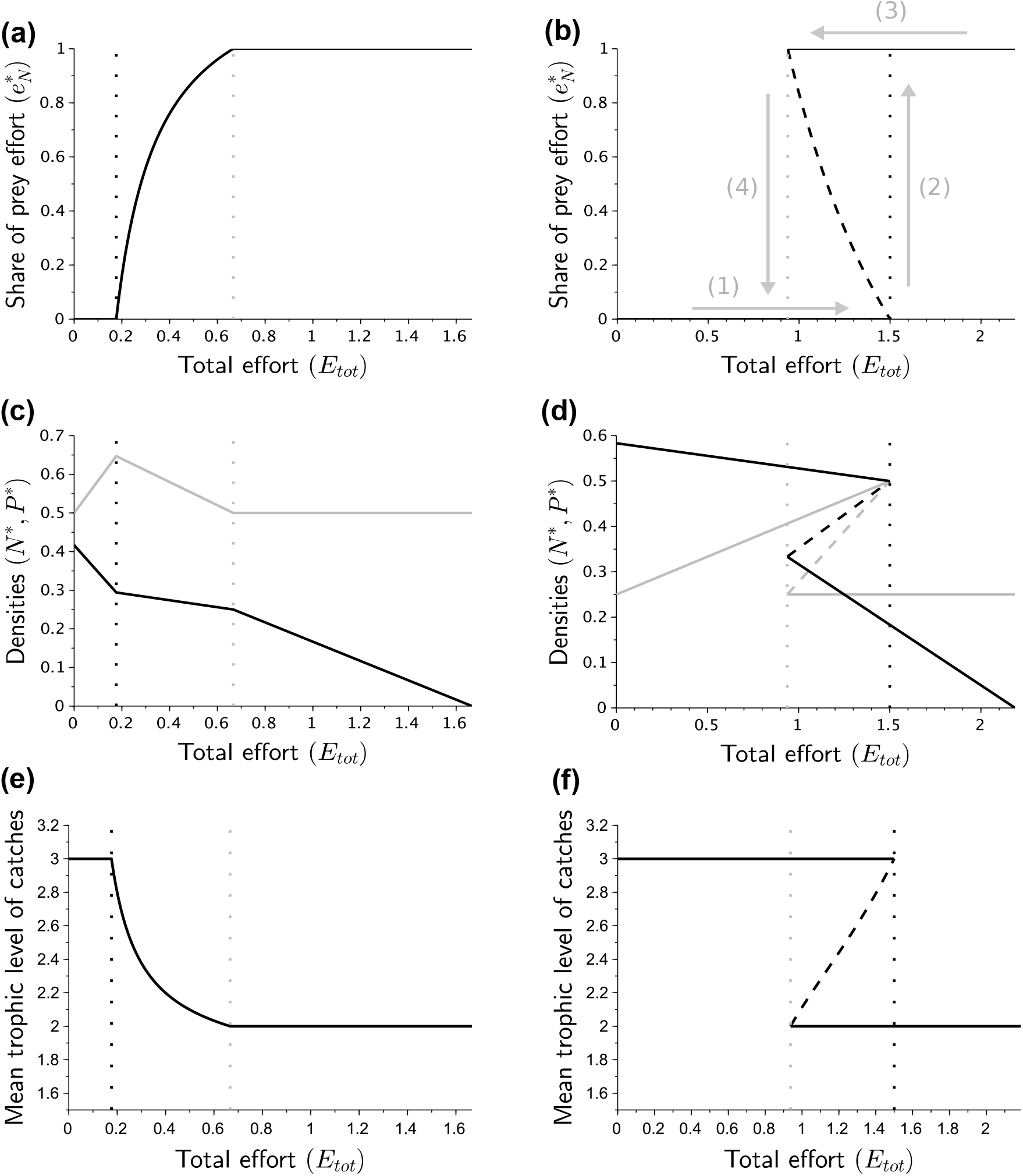
Share of prey effort (a,b), equilibrium densities (c,d) and mean trophic level of catches (e,f) for increasing fishing efforts. When condition (12) is verified (a,c,e), the mixed equilibrium is stable, and when it is not verified (b,d,f), the system exhibits alternative stable states. (c,d) gray line: prey density; black line: predator density. Dotted lines represent 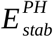 (black dotted line) and 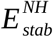 (gray dotted line). Dashed lines indicate unstable equilibria. In (b), arrows are used to describe the phenomenon of *hysteresis* (see the description of the different steps labeled (1) to (4), in the text). Parameters in (a,c,e): *r*=1, *K*=1, *m*=0.3, *μ*=0.5, *γ*=1.2, *q*_*N*_=0.3, *q*_*P*_=0.5, *p*_*N*_=1, *p*_*P*_=2, *c*_*N*_=0.1, *c_P_*=0.2. Parameters in (b,d,f): same as (a,c,e) except *K*=2, *μ*=0.8, *γ*=1.5, *q*_*N*_=0.4, *q*_*P*_=0.2 and *p*_*P*_=3,.

Sudden regime shifts between the predator- and the prey-focused equilibria occur whenever the mixed harvest stability condition - condition (12) - does not hold. In that case, increasing the total fishing effort induces a sudden shift from a predator- to a prey-focused fishery, characterized by a sudden decrease in the mean trophic level of catches (Fig. 1f). When the mixed harvest stability condition - condition (12) - does not hold, the effort below which the predator-focused system is stable 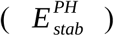 is larger than the effort above which the prey-focused system is stable 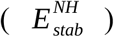. In between, the predator- and the prey-focused equilibria are simultaneously stable, while the mixed harvest equilibrium is unstable (see Fig. 1b). This bistable system is characterized by a phenomenon called *hysteresis*: starting from a predator-focused system, increasing the fishing effort (arrow (1) in Fig. 1b) leads to a regime shift towards a prey-focused system when the effort reaches 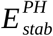 (arrow (2) in Fig. 1b), inducing severe losses in both prey and predator densities (Fig. 1d); then, to recover higher prey and predator densities, managers would have to reduce the fishing effort to 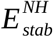 (arrows (3) and (4) in Fig 1b).

To better understand the mechanism behind these regime shift dynamics, we study the different feedback loops that stabilize or destabilize our system. To do this we consider the non-diagonal elements of the Jacobian matrix (see Appendix A), which contains partial derivatives of our model, evaluated at the equilibrium. It informs us about the sign of the relationship between two variables, and about the parameters that are involved in this relationship. We show in Figure 2 the relationships between prey density, predator density, and share of prey effort. We also indicate the parameters involved both in the strength of the interaction and in the mixed harvest stability condition - condition (12).

**Figure 2:**
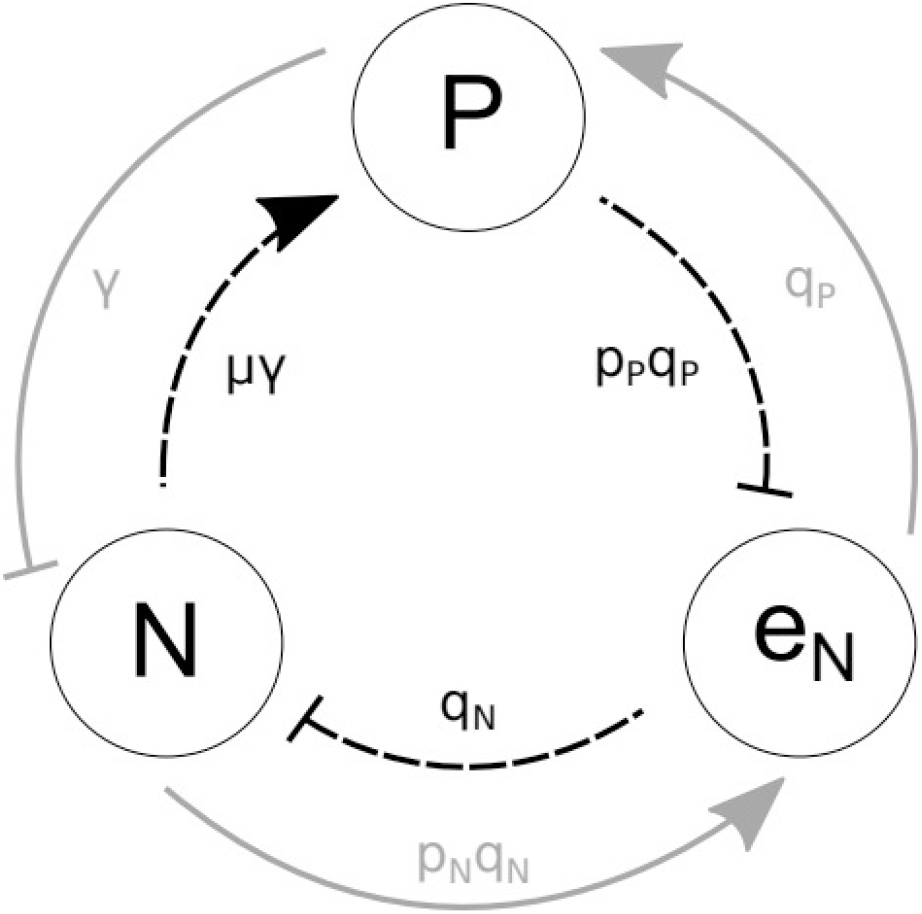
Illustration of the relationships between predator density, prey density and share of prey effort. Arrows indicate positive relationships, while bars indicate negative relationships. Gray lines form the stabilizing loop, and black dashed lines form the destabilizing loop. Parameters *p*_*N*_ and *p_P_* are the respective prices of prey and predators, *γ* the predator attack rate, *μ* the prey-to-predator conversion efficiency, *q_N_* and *q_P_* the prey and predator catchabilities.

Two loops are involved: the first entails one negative feedback and is therefore stabilizing (gray lines); the second loop creates a global positive feedback and is therefore destabilizing (dashed black lines). Suppose that there is an increase in the share of prey effort. Following the gray loop, this reduces the fishing pressure on predators, as modulated by catchability *q*_*P*_, therefore increasing predator density. This leads to increased top-down controls, mediated by attack rate *γ*, thus reducing prey density. This decreases the profitability of prey harvesting, as constrained by parameters *p_N_* and *q*_*N*_, thereby reducing the share of prey harvest. This first loop is therefore stabilizing. But following the black loop, the increase in the share of prey effort enhances the fishing pressure on prey, modulated by catchability *q*_*N*_, thereby decreasing prey density. This induces a negative bottom-up effect mediated by parameters *μ* and *γ*, that reduces predator density. This decreases the profitability of predator harvesting, as constrained by parameters *p_P_* and *q*_*P*_, thus further increasing the share of prey harvest. This second loop is therefore destabilizing.

The relative influence of the stabilizing or of the destabilizing loop is dependent on the value of parameters involved in the mixed harvest stability condition (12), that determine the stability of the mixed harvest equilibrium. For instance, a low predator price and a high prey price favor the stabilizing loop. On the contrary, a high conversion efficiency favors the destabilizing loop. The roles of the other parameters involved in the mixed harvest stability condition (12) are dependent on the value of (*μ* − *p_N_* / *p_P_*), which explains why they are involved in both stabilizing and destabilizing loops. For instance, increasing conversion efficiency *μ* promotes the stabilizing effect of predator catchability *q*_*P*_, while increasing the prey-to-predator price ratio *p*_*N*_ / *p_P_* enhances the destabilizing effect of predator catchability.

We now investigate the impact of adaptation speed *G* on the resilience of the system. Let us assume that the mixed harvest stability condition (12) holds, which implies that a mixed harvest is possible. As illustrated in Figure 3a, increasing the pace of adaptation is first stabilizing in a predator- and in a prey-focused fishery until resilience reaches a plateau. This pattern can also be inferred from the analytical expressions of the eigenvalues (see Appendix A). In a mixed fishery, while low to intermediate adaptation speeds stabilize the system, higher adaptation rates turn out to be destabilizing.

**Figure 3:**
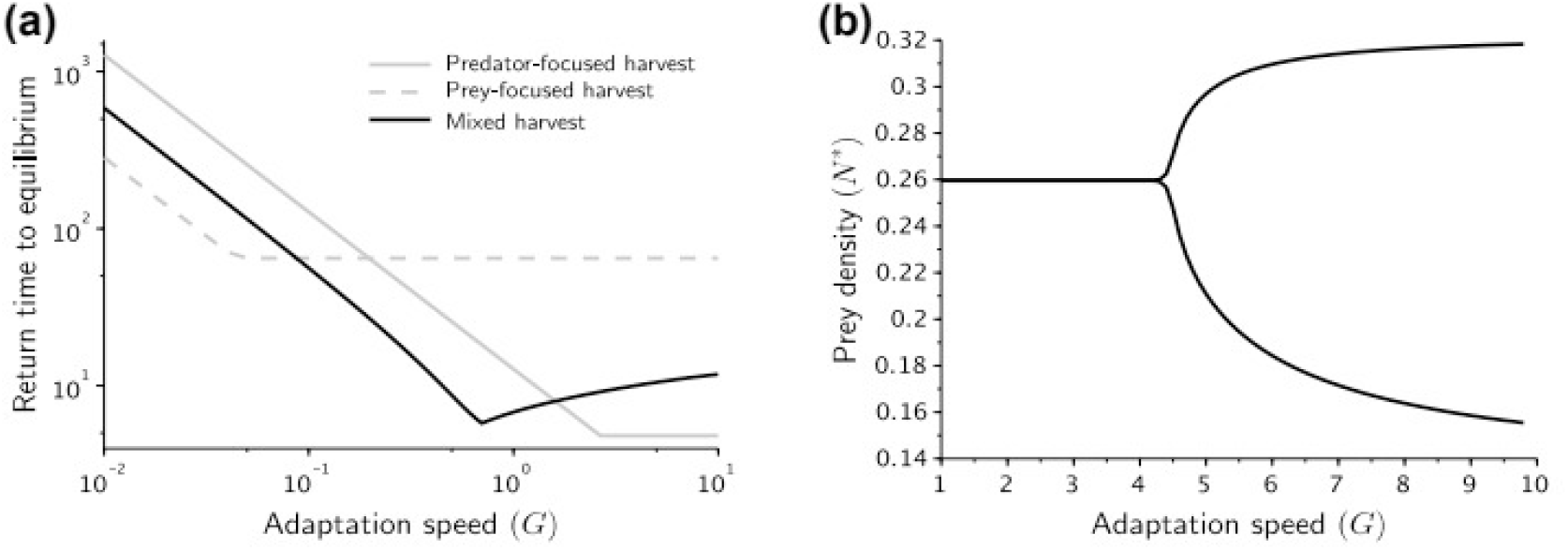
Influence of adaptation speed on stability. (a) Relationship between return time to equilibrium and adaptation speed in the predator-focused (*E*_*tot*_=0.3), the prey-focused (*E*_*tot*_=2.7) and the mixed equilibrium (*E*_*tot*_=1). Parameters: *r*=1, *K*=1, *m*=0.1, *μ*=0.5, *γ*=1.2, *q*_*N*_=0.3, *q*_*P*_=0.5, *p*_*N*_=1, *p*_*P*_=2, *c*_*N*_=0.1, *c_P_*=0.2. (b) Minimum and maximum prey densities for increasing adaptation speeds in a Rosenzweig-MacArthur model at the mixed equilibrium. Parameters: same as (a), except *K*=1.5, *m*=0.2, *N*_0_=0.4, *E*_*tot*_=1.75.

Because return time to equilibrium is only one aspect of resilience, we also study how adaptation speed affects population variability. To do so, we study a predator-prey model with a Holling type-II functional response, described in equation (3). As analytical work is made difficult by the non-linear functional response, we run simulations of 50000 time steps for different values of adaptation speed. We choose parameters so as to be in a mixed harvest situation, where 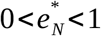. Results are shown in Fig. 3b. For low adaptation rates, the system is stable and does not oscillate. But higher adaptation rates induce oscillations, with growing amplitude. In addition to increasing the return time to equilibrium, high adaptation speeds therefore drive the system into an unstable oscillating state. In Appendix B, we show that the negative impact of adaptation speed on stability is robust to changes in parameters.

The model we present here is simplified, focusing on the effects of adaptive fishing on two species. This allowed us mathematically analyze the system and to show that fishing down the web patterns occur in all instances, due to effort reallocation based on species profitabilities. A legitimate question is whether our main results hold in more complex settings. To investigate this, we extended adaptive harvesting to three species systems.

### Three-species fishery

We numerically investigate the effects of increasing the total fishing effort on the behavior of the three-species model. According to analytical results from the predator-prey model, we hypothesize a fishing down effect under increasing efforts, that is a shift from a fishery focused on the highest trophic level to a fishery focused on the lowest trophic level, with a transitory mixed fishery where the effort is shared between species.

We first investigate the adaptive harvest of a trophic chain, where there is no direct link between the predator and the resource 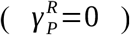. Results are shown in Figure 4. For very low fishing efforts, the fishery only focuses on the predator. Increasing the fishing effort therefore reduces predator density, which in turn increases consumer density and decreases resource density. As consumers become more abundant, their relative marginal profitability increases, leading to a first shift from a predator-focused fishery to a consumer-and-predator mixed fishery. In this fishery, as both predator and consumer are harvested, increasing fishing efforts lead to reduced consumer and predator densities, and to increased resource density. Reduced predator densities bring a second shift towards a fishery where only the consumer is harvested. In this consumer-focused fishery, consumer and resource densities remain stable, while predator densities decrease towards extinction. After this extinction, increasing the fishing effort leads to a third shift towards a mixed consumer-and-resource fishery. Increasing fishing efforts in this mixed fishery reduces the densities of both harvested species, until they both get extinct. In parallel, the fishery gradually shifts towards a resource-only fishery.

**Figure 4:**
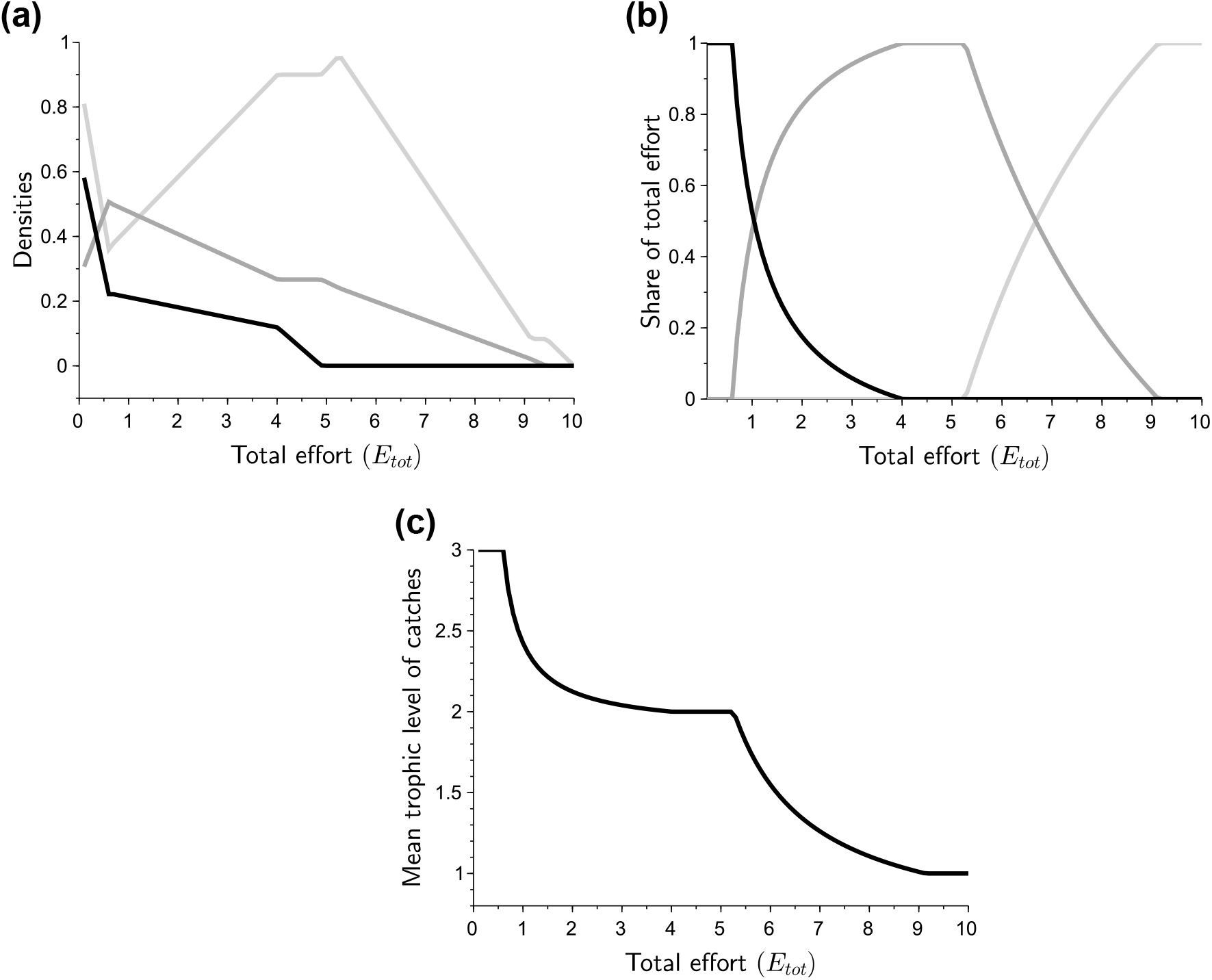
Equilibrium densities (a), share of total effort (b), and mean trophic level of catches (c) for increasing fishing efforts in a tritrophic food chain. (a) light gray line: resource density; dark gray line: consumer density; black line: predator density. (b) light gray line: share of resource effort; dark gray line: share of consumer effort; black line: share of predator effort. Value of fixed parameters: *r*=1, *K*=1.5, *m_C_*=0.1, *m_P_*=0.2, 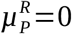, 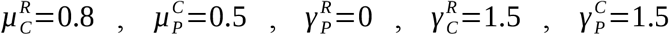, *q_R_*=0.1, *q_C_*=0.2, *q_P_*=0.3, *p*_*R*_=1, *p_C_*=2, *p*_*P*_=3, *c*_*R*_=0.1, *c_C_*=0.1, *c*_*P*_=0.1, *G*=1.

Impacts of increasing total fishing effort on the mean trophic level of catches are shown in Figure 4.c. We observe a globally declining mean trophic level of catches with increasing fishing efforts, which is consistent with our prediction from the predator-prey model. As higher trophic levels are gradually depleted with increasing fishing efforts, the trophic level of the community also undergoes a general decrease.

Additionally, we investigate increasing fishing efforts in a tritrophic food web displaying omnivory 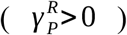. Results are shown in Figure 5. Results are globally coherent with the food chain simulations : increasing fishing efforts induces a shift from a predator-focused fishery to a resource-focused fishery, which is characteristic of a fishing down the food web pattern. While the fishing down the web pattern therefore again emerge from the adaptive harvesting, we note that, interestingly, a mixed fishery where all three species are harvested is selected at intermediate effort in omnivory modules. In this fishery, increasing the fishing pressure jointly reduces the densities of all three species, until predator densities become too small to be interesting to harvest. Increasing fishing efforts indirectly reduces predator densities, while keeping consumer and resource densities stable. As reduced predator densities release pressure on consumers, the share of effort dedicated to consumers increases, leading to a very slight increase in the mean trophic level of catches (when E_tot_ is a little below 8 on Fig 5c). This very slight « fishing up » the food web does not however alter the global conclusion, as fishing down the web patterns remain largely dominant (Fig 5c) can also emerge in developing fisheries. After predator extinction, increasing the fishing effort brings an increased focus on resource, leading to a decreased mean trophic level of catches until both consumer and resources get extinct. We therefore show that the fishing down the food web pattern we observed in the two species model (main text), also emerge from the adaptive harvesting of a tri-trophic food web.

**Figure 5:**
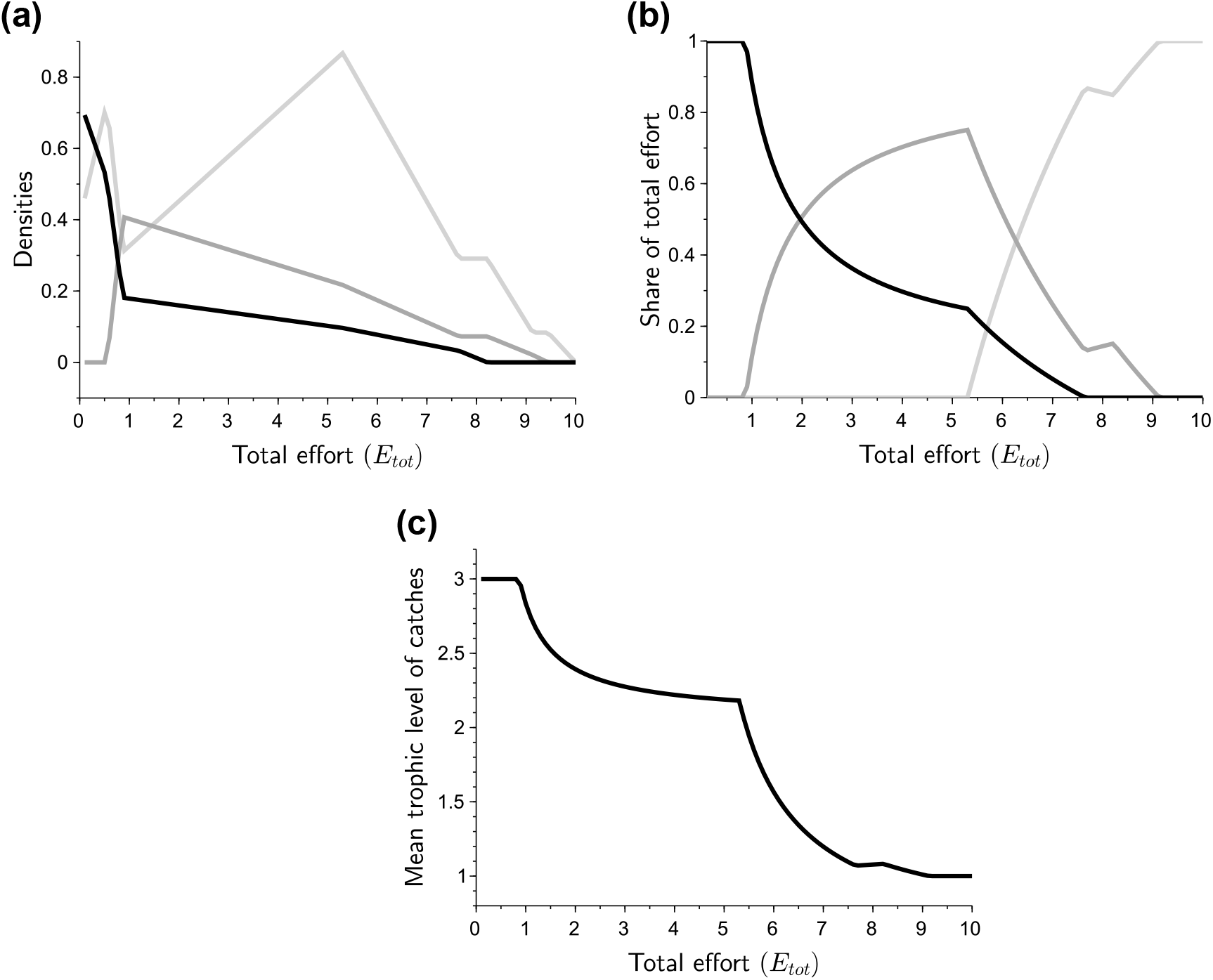
Equilibrium densities (a), share of total effort (b), and mean trophic level of catches (c) for increasing fishing efforts in a tritrophic food web with omnivory. (a) light gray line: resource density; dark gray line: consumer density; black line: predator density. (b) light gray line: share of resource effort; dark gray line: share of consumer effort; black line: share of predator effort. Value of fixed parameters: *r*=1, *K*=1.5, *m_C_*=0.1, *m*_*P*_=0.2, 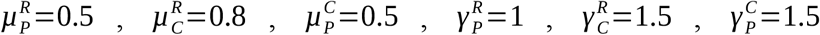, *q_R_*=0.1, *q*_*C*_=0.2, *q_P_*=0.3, *p*_*R*_=1, *p_C_*=2, *p*_*P*_=3, *c*_*R*_=0.1, *c_C_*=0.1, *c*_*P*_=0.1, *G*=1.

Finally, we assess the possibility of regime shifts in tritrophic communities. With the two-species model, we show that bistable systems can emerge, notably when top-down effects are large, or when the prey-to-predator price ratio is low. We show in Figure 6 that such bistable systems also emerge under similar conditions in a tritrophic food chain (Fig. 6a) and in a tritrophic food web with omnivory (Fig. 6b). To do so, we use sets of parameters that are similar to those inducing bistability in the predator-prey system (Figure 1b and 1d). As we increase the fishing effort, we observe in both cases a progressive shift from a predator-focused to a mixed predator-consumer fishery, and then a sudden shift from a mixed predator-consumer fishery to a mixed predator-resource fishery, that translates into a sudden drop in the mean trophic level of catches (indicated by arrows in Figure 6). Inversely, when the total fishing effort decreases, we observe a sudden increase in the mean trophic level of catches. The upwards shift occurs at a lower effort than the downwards shift, a phenomenon called *hysteresis*, that is characteristic of bistable systems.

**Figure 6:**
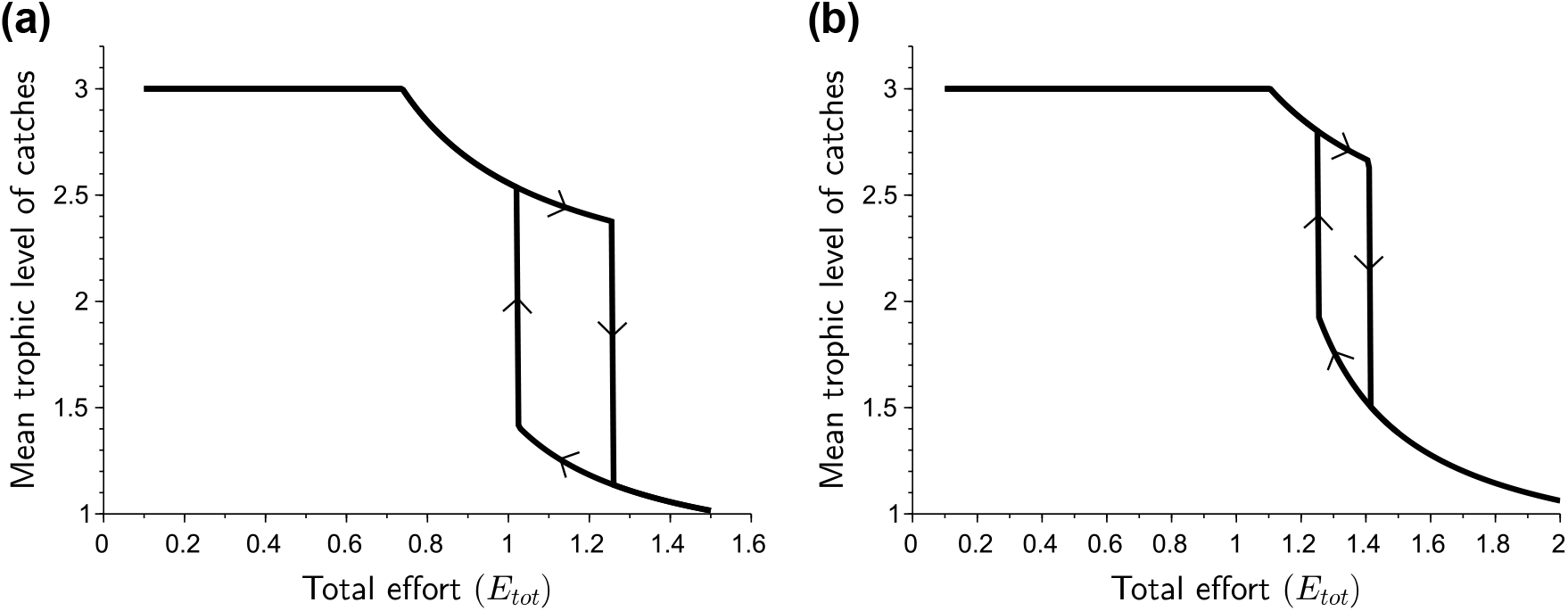
Mean trophic level of catches for increasing fishing efforts in a tritrophic food chain (a) and in a tritrophic food web with omnivory (b). (a) Value of fixed parameters: *r*=1, *K*=2, *m_C_*=0.3, *m_P_*=0.1, 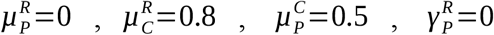, 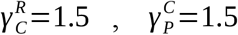, *q_R_*=0.4, *q_C_*=0.2, *q_P_*=0.4, *p_C_*=1, *p_C_*=2, *p_P_*=4, *c*_*R*_=0.1, *c*_*C*_=0.2, *c_P_*=0.1, *G*=1. (b) Same as (a), except 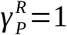 and 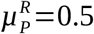.

This additional result suggests that sudden regime shifts also occur in more complex adaptively harvested communities. Further numerical work could help understand stability conditions in such systems.

## Discussion

Our results suggest that fishing down the food web - associated with a decline in the mean trophic level of catches and in the mean trophic level of the ecosystem - can result from adaptive foraging by fishermen. This view is already contained in the original fishing down theory (Pauly et al. 1998), but subsequent studies mainly interpreted lower trophic levels of catches as resulting only from overfishing and declining predator abundances, without accounting for the fact that fishermen can re-allocate their fishing effort to lower trophic levels (Essington et al. 2006).

Theoretical studies support the view that fishing down processes emerges from feedbacks between changing fish abundances and adaptive fishing patterns. Branch et al. (2010) found for instance that adaptive fishing patterns based on stock availability reduce the mean trophic level of catches in ecosystem models. Andersen et al. (2015) also showed with a size-spectrum fishery model that increasing the global fishing pressure leads to a gradual re-allocation of effort from big predatory fish to medium and small prey fish. Likewise, Wilen and Wilen (2012) found that if top predators in a food chain model have high values, then profit-driven effort dynamics first imply a focus on the top trophic level, followed by an increase in efforts targeting lower trophic levels. Similar results have been obtained with an age-structured model of a food chain (Wiedenmann et al. 2016).

Furthermore, empirical evidence suggest shifts in target from upper- to lower-trophic-levels in various fisheries. In the Scotian Shelf fishery, the decline in the groundfish community caused a trophic cascade that produced a new fishing regime, dominated by shrimp and crab landings (Frank et al. 2005). Likewise, in the Argentinean-Uruguayan common fishing zone, data suggest that the fishing effort has been redistributed from overexploited to more abundant species (Jaureguizar and Milessi 2008). However, precise empirical investigations of this ecological-economic feedback are lacking.

Originally, fishing down the food web has been described using the mean trophic level of catches as an indicator (Pauly et al. 1998). Branch et al. (2010) showed however that trends in this indicator can diverge from trends in the mean trophic level of the ecosystem. This can be explained by the fact that the mean trophic level of the ecosystem only reflect changes in abundances, while the mean trophic level of catches also reflects changes in fishing strategies (shifting from lower to higher trophic levels). As in the present paper we are interested in how fishing strategies change with increasing fishing effort, we chose to focus on the original mean trophic level of catches indicator.

While adaptation is here in the harvesting regime, some of our results are coherent with theoretical work on adaptive foraging of predators in ecological models. (Křivan and Schmitz 2003) for instance studied optimal predator foraging on a consumer-resource community. They show that at low densities, the top predator focuses on the consumer, at higher densities it focuses on the resource, and at intermediate densities it predates on both consumer and resource. This is very similar to the relationship we find between the total fishing effort and the fishing pattern. The main difference is that while natural predators are expected to maximize energy inputs, in our model fishermen’s behavior aims at maximizing profits. Moreover, our model allows to account for changes in adaptation speeds, while in (Křivan and Schmitz 2003) adaptation is instantaneous.

Adaptive foraging is often thought to be stabilizing (Křivan 2007). Particularly it facilitates prey coexistence, as predation is relaxed when prey becomes rare (Kondoh 2003). Adaptive foraging of predators may however alter the intensity of apparent competition by changing predator density, leading to the emergence of instabilities (Krivan 1996). An important difference is that in the present model, consumption pressure is exerted by the fishing effort, that does not have any population dynamics. We find that adaptive harvesting can then induce sudden regime shifts. More generally, as pointed out in (Lade et al. 2013), social-ecological feedbacks may lead to regime shifts and bistability, while similar uncoupled systems do not. Using a harvested predator-prey model, (Horan et al. 2011) also found that modifying the feedback response between populations and management alters the stability landscape of the system. However, many studies on ecological regime shifts focus on ecological drivers and consequences of these shifts (Conversi et al. 2014). Reconsidering the role of human dynamics in triggering regime shifts in marine food webs could thus help understand and prevent such phenomena.

Our investigation suggests that large top-down effects vs. bottom-up effects favor regime shifts. It thus echoes studies stressing the importance of top-down forcing for triggering regime shifts in marine food webs (Pershing et al. 2014). Moreover, we show that the possibility of a social-ecological regime shift is dependent on the relative intensity of a stabilizing and a destabilizing feedback loop between prey and predator densities and the share of prey harvest. We notably find that high conversion efficiencies promote the destabilizing loop, while a high prey-to-predator price ratio enhances the stabilizing loop. This stresses the joint influence of ecological and economic drivers in triggering regime shifts, as pointed out in (Lade et al. 2015).

The propensity of undergoing regime shifts is only one of the many facets of the resilience of a system (Donohue et al. 2016). We also investigated the impact of adaptation speed on the ability of a system to recover from a perturbation, and on the speed of this recovery. In a prey- and a predator-focused equilibrium, adaptation speed is always stabilizing, as it reduces return times to equilibrium. In a mixed fishery, only low adaptation speeds are stabilizing. This is quite important as we expect that adaptation may be quite slow in many instances, for instance due to technological or regulation constraints. Therefore, the fact that many regulated fisheries seem to be sustainable given recent data (Duarte et al. 2020) would fit such situations of stable, mixed harvesting situations that we expect for intermediate (regulated) effort and low adaptability. Our results are also coherent with former studies. Bischi et al. (2013) found with a harvested predator-prey model with switching fishing targets, that the speed of convergence is higher and that oscillations are reduced when the switching time approaches zero. It is also coherent with studies showing that predator’s adaptation speed increases the resilience of food webs (Kondoh 2003).

High adaptation speeds however reduce the resilience of mixed fisheries, which contradicts our initial expectation of a stabilizing adaptive foraging. We find that high adaptation speeds increase the return time to equilibrium in a linear predator-prey system, and induce oscillations in a non-linear predator-prey model. In the latter case, the amplitude of oscillations increases with adaptation speed. Likewise, (Kramer 2008) showed that a theoretical harvested reef ecosystem displays larger variations with high adaptation speeds. (Bischi et al. 2013) also showed with a harvested predator-prey model that fast switching from predator to prey induces oscillatory behaviors. Similarly, using an age-structured food chain model, (Wiedenmann et al. 2016) found that biomass, catches and profits are more variable when fishermen adaptively switch their target. (Abrams 1999) also showed that adaptive foraging of a predator on two prey could yield unstable dynamics, and that these dynamics are dependent on the rate of predator adaptation.

Social-ecological resilience therefore depends on how fast fishermen or management authorities react to changes in relative abundances and profits. We have voluntarily kept the model simple and generic, to discuss the effects of adaptive fisheries at different scales of organization. Our model may for instance be construed at the scale of individual fishermen. At this scale, agents may imitate the most successful strategy or change their strategy based on previous success so that adaptation speed may be quite rapid (Noailly et al. 2003). However, we expect fast changes to only occur when access to the different species is not strongly regulated (eg, open access fisheries), and when such changes are not constrained by changes in equipment or technologies (Noailly et al. 2003, Bischi et al. 2013). Our model can also be used to understand changes in regulation at higher organizational scales, for instance changes in regulation of the effort among species by authorities. We then expect delays inherent to collective decision-making and therefore expect slow adaptation. Note however that harvesting strategies and policies rely on frequent population census and profitability assessments, and are thus adaptive. We therefore expect that our model may provide a good basis to understand such changes.

Including such adaptive dynamics in management models and plans is critical to achieve a sound management of multispecies fisheries. In particular, our results suggest that such adaptation may severely constrain different aspects of fisheries resilience. We therefore suggest that considering adaptive economic behaviors is a key step towards the implementation of an ecosystem-based fisheries management (Pikitch et al. 2004). While our results seem to be robust to changes in the community context, as fishing down the web patterns emerge from adaptive harvesting in all two species and three species modules we considered, the models used here are still very simple. Particularly, as often done in theoretical ecology, we analyse the stability of equilibria of deterministic models. In real situations, external disturbances occur, both in the environmental conditions and in the economic scenarios. Multiple disturbance scenarios could be considered and they may affect resilience and sustainability in different ways. Recent papers have also highlighted that the impact of fishing low trophic level forage fish on their predators is often less than expected using usual trophic models (Hilborn et al. 2017), and that predators exposed to high fishing mortality can actually benefit from harvesting of their forage fish (Soudijn et al. 2021). We believe understanding adaptive harvesting including in these more complex situations may open new doors for a successful management of harvested ecosystems.

## Appendix A: Analytical calculations

We here describe the three possible equilibria where predator and prey coexist, with their associated feasibility and stability conditions. These equilibria are illustrated in Figure A1, and their analytical expressions can be found in the following table:

**Table A1:** Description of the three considered equilibria.

| Predator harvest (PH) | Prey harvest (NH) | Mixed harvest (MH) |
| --- | --- | --- |
| $N^* = \frac{m + q_P E_{tot}}{\mu \gamma}$ $P^* = \frac{r}{\gamma} \left( 1 - \frac{N^*}{K} \right)$ $e_N^* = 0$ | $N^* = \frac{m}{\mu \gamma}$ $P^* = \frac{1}{\gamma} \left[ r \left( 1 - \frac{m}{\mu \gamma K} \right) - q_N E_{tot} \right]$ $e_N^* = 1$ | $N^* = \frac{m + q_P (1 - e_N^*) E_{tot}}{\mu \gamma}$ $P^* = \frac{p_N q_N N^* + c_P - c_N}{p_P q_P}$ $e_N^* = \frac{1}{q_N E_{tot}} \left[ r \left( 1 - \frac{N^*}{K} \right) - \gamma P^* \right]$ |

**Figure A1:**
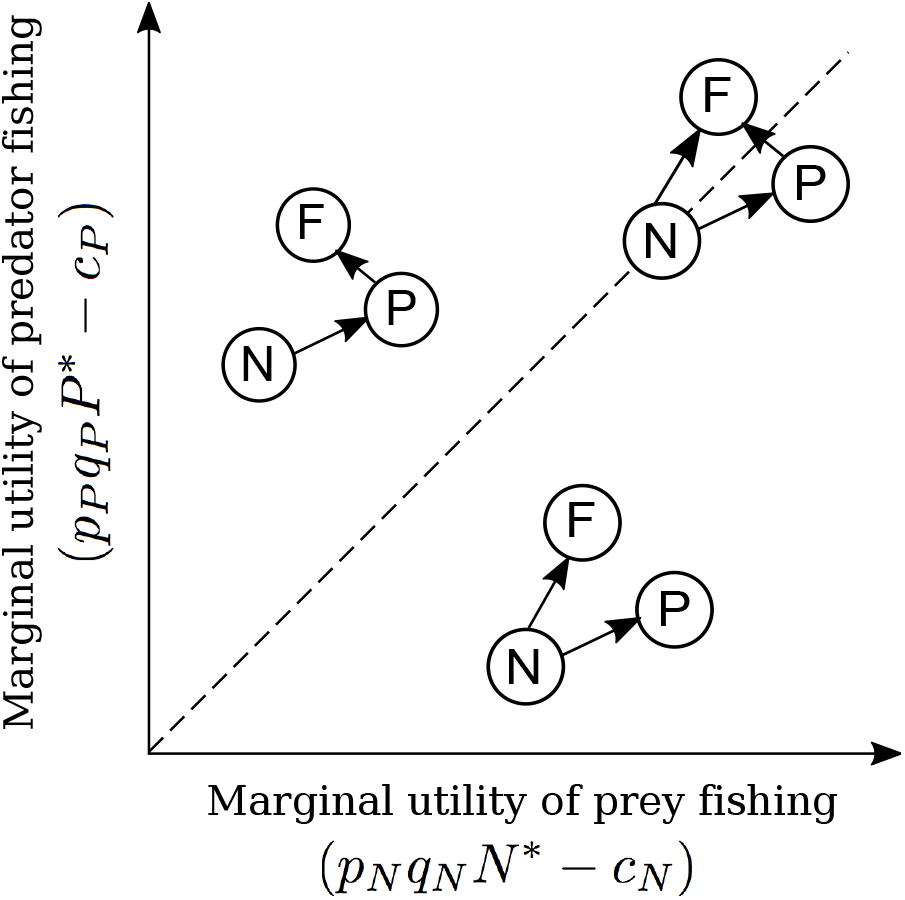
Illustration of the different fishing patterns that emerge from adaptive harvesting. Fishermen are described by the letter F, prey by N and predators by P. If the marginal utility of harvesting the predator is larger than the marginal utility of harvesting the prey, then the harvest is focused on predators. If it is smaller, then the harvest is focused on prey. If the marginal utilities of harvesting prey and predators are equal, then the fishing effort is shared between the two species.

An equilibrium is feasible for parameter sets that ensure that prey and predator densities are positive. The mixed equilibrium is characterized by a supplementary feasibility condition, namely that the share of prey effort is between 0 and 1. We further assume that coexistence of prey and predator in the unharvested system is warranted, which implies that *m*<*μγK*. The local stability of the system can be assessed by computing the eigenvalues of the Jacobian of the system at equilibrium. The general formulation of this Jacobian is:

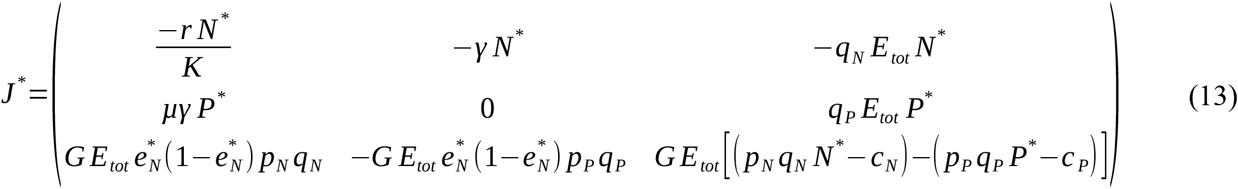

### 1. Predator-focused harvest (PH)

#### 1.1. Feasibility conditions

Prey density *N*^*^ is always positive and increases with the fishing effort. Predator density *P*^*^ is positive when

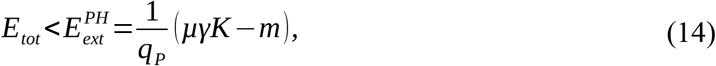

and it decreases with the fishing effort. The effort below which the predator-focused harvest is feasible is denoted 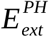.

#### 1.2. Stability conditions

The eigenvalues of the system can be written:

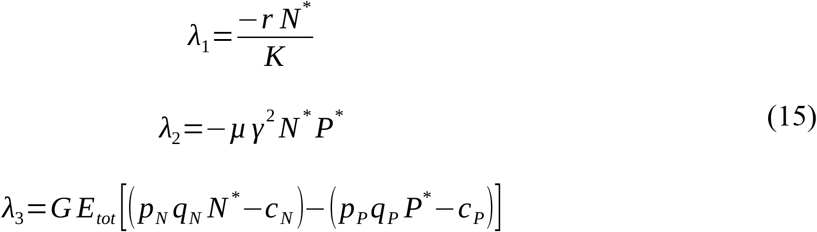

Eigenvalues *λ*_1_ and *λ*_2_ are negative if the feasibility conditions are fulfilled. The third eigenvalue *λ*_3_ is negative as long as *p_P_q_P_P*^*^−*c_P_* > *p_N_q*_*N*_*N*^*^−*c_N_*. This translates into the following condition:

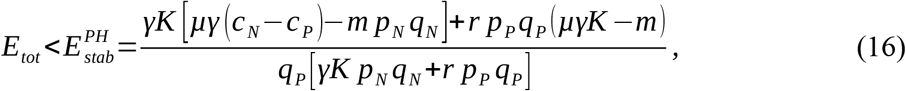

where the effort below which the predator-focused harvest is stable is denoted 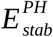. All feasible states are not stable if 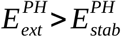, which translates into the following condition:

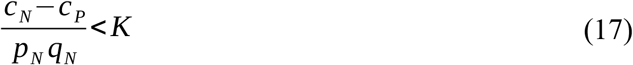

This condition always holds if the cost of harvesting the prey is smaller than the cost of harvesting the predator (*c*_*N*_ < *c*_*P*_).

### 2. Prey-focused harvest (NH)

#### 2.1. Feasibility conditions

Prey density *N*^*^ is always positive. Predator density *P*^*^ is positive when

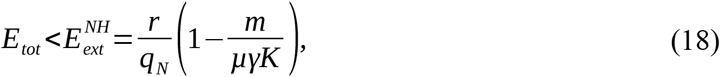

and it decreases with the fishing effort. The effort below which the prey-oriented harvest is feasible is denoted 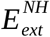.

#### 2.2. Stability conditions

The eigenvalues of the system can be written:

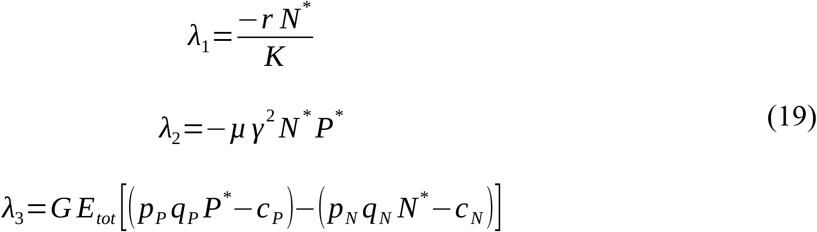

Eigenvalues *λ*_1_ and *λ*_2_ are negative if the feasibility conditions are fulfilled. The third eigenvalue *λ*_3_ is negative as long as *p_N_q_N_N*^*^−*c_N_* > *p_P_q*_*P*_*P*^*^−*c_P_*. This translates into the following condition:

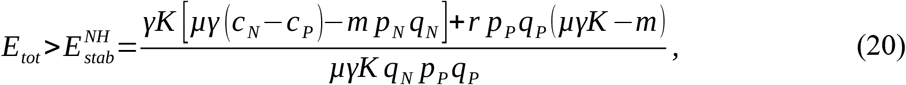

where 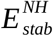 stands for the effort above which the prey-focused harvest is stable. If 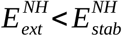, none of the feasible states are stable. On the contrary, some feasible states can be stable if 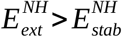, which translates into the following condition:

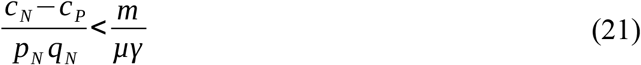

If the harvesting the predator is costlier than harvesting the prey (*c*_*N*_ < *c*_*P*_), this condition always holds.

### 3. Mixed predator-prey harvest (MH)

#### 3.1. Feasibility conditions

As 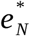 is supposed to take values between 0 and 1, *N*^*^ is always positive. We can specify the effort at which the predator goes extinct by using the complete expression of the predator density at equilibrium:

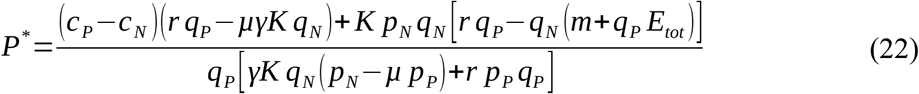

Predator densities are positive if:

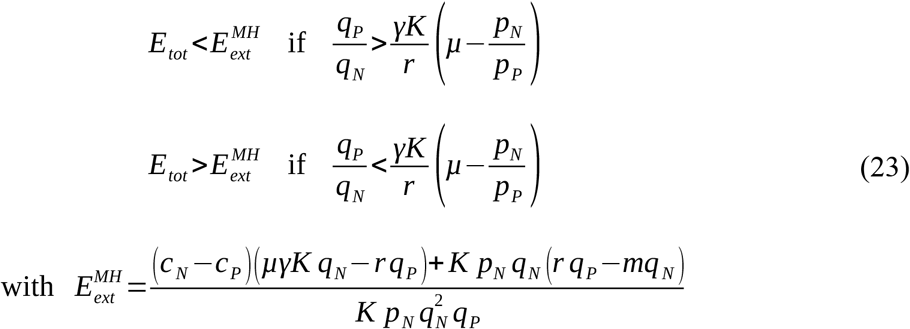

Here with 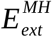 stands for the effort at which predators goes extinct in the mixed harvest equilibrium. Finally, as the share of prey effort can be written

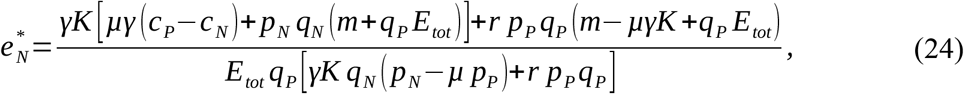

the feasibility condition on 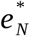 becomes:

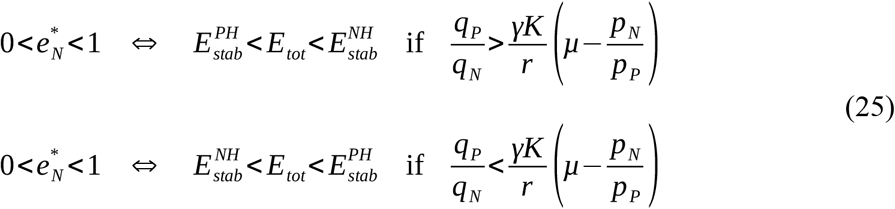

Note that the second case is only possible if *p*_*N*_ / *p_P_* < *μ*, that is if the ratio of prey to predator prices is smaller than the prey-to-predator conversion efficiency.

#### 3.2. Stability conditions

The analysis of stability can be done by computing the Jacobian matrix and the associated Routh-Hurwitz conditions. The characteristic polynomial *P*(*λ*)=*a_0_λ*^3^+*a_1_λ*^2^+*a_2_λ*^1^+*a_3_* can be written as follows:

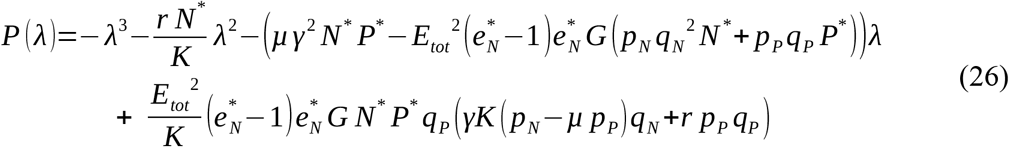

The system is stable provided *a*_0_, *a*_1_, *a*_2_, *a*_3_<0 and *a*_1_ *a*_2_−*a*_0_ *a*_3_>0. The coefficient *a*_0_, *a*_1_ and *a*_2_ are negative as long as the feasibility conditions are verified. The coefficient *a*_3_ is negative if

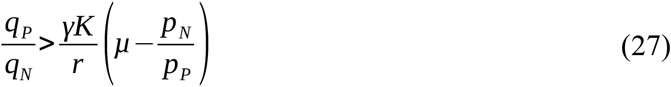

The stability condition *a*_1_ *a*_2_−*a*_0_ *a*_3_>0 is equivalent to

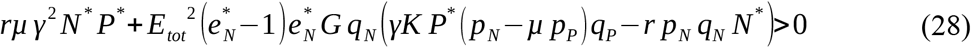

If *p_N_* ≤ *μp_P_*, this condition is always verified, and stability only depends on condition (27). If *p*_*N*_ >*μp_P_* however, condition (27) is always true and stability then depends solely on condition (28), which is verified if

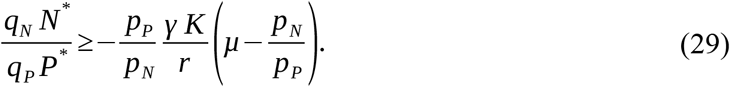

From condition (27), this implies that

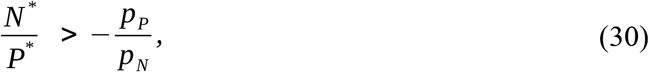

which is always true as *N*^*^ and *P*^*^ are positive. In short, if *p_N_*>*μp_P_*, then the equilibrium is stable, and if *p_N_* ≤ *μp_P_*, then the stability of the equilibrium depends on condition (27). As shown in the main text, if condition (27) is not true, the system is bistable.

#### 3.3. Effect of fishing effort on densities

Let us now investigate the precise effect of fishing pressure on the prey and predator densities. The following relationship can be established:

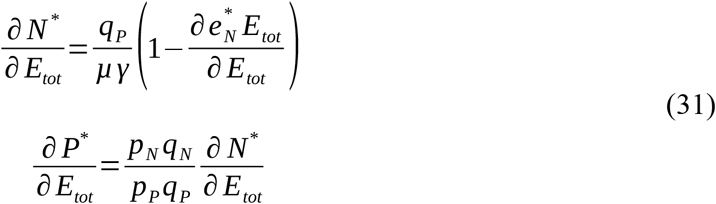

Prey and predator densities thus vary in the same direction with increasing efforts. This direction is dependent on the effect of effort on 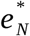. As we have

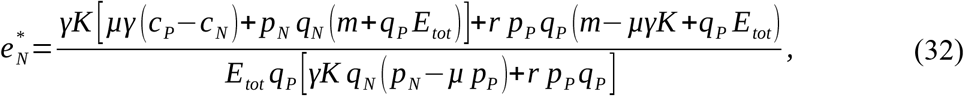

we get the following derivative:

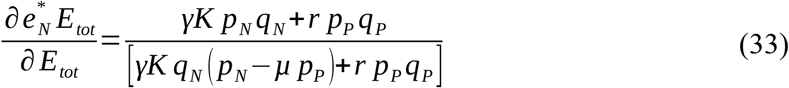

Then, two different cases appear:

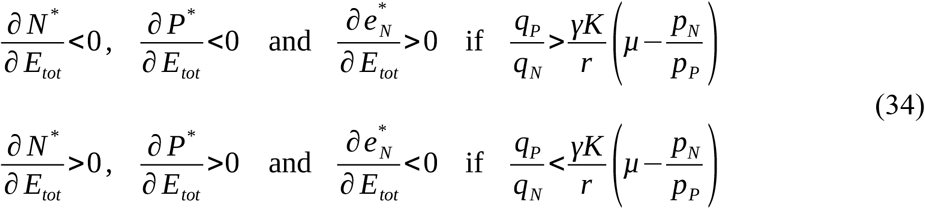

Thus, if the system is stable, predator and prey densities decrease while the share of prey harvest increases with the fishing effort, and if the system is bistable, predator and prey densities increase while the share of prey harvest decreases with the fishing effort.

### 4. Summary table

In the following table, we summarize the feasibility and stability conditions of the different possible equilibria. This helps to understand the relationships between these equilibria.

**Table A2.** Feasibility and stability conditions of the different equilibria. Expressions of the limit efforts are found in the Appendix. Only stability conditions that are not redundant with feasibility conditions are shown.

| Equilibrium | Predator harvest<br>(PH) | Mixed harvest (MH) | Prey harvest<br>(NH) |
| --- | --- | --- | --- |
| Feasibility<br>conditions | $E_{tot} < E_{ext}^{PH}$ | $\text{if } \frac{q_P}{q_N} > \frac{\gamma K}{r} \left( \mu - \frac{p_N}{p_P} \right):$ $E_{tot} < E_{ext}^{MH}$ $E_{stab}^{PH} < E_{tot} < E_{stab}^{NH}$ | $E_{tot} < E_{ext}^{NH}$ |
| | | $\text{if } \frac{q_P}{q_N} < \frac{\gamma K}{r} \left( \mu - \frac{p_N}{p_P} \right):$ $E_{tot} > E_{ext}^{MH}$ $E_{stab}^{NH} < E_{tot} < E_{stab}^{PH}$ | |
| Stability<br>conditions | $E_{tot} < E_{stab}^{PH}$ | $\frac{q_P}{q_N} > \frac{\gamma K}{r} \left( \mu - \frac{p_N}{p_P} \right)$ | $E_{tot} > E_{stab}^{NH}$ |
| Relationships<br>between<br>conditions | $E_{ext}^{PH} > E_{stab}^{PH}$ ⇔ $\frac{c_N - c_P}{p_N q_N} < K$ | $\text{if } \frac{q_P}{q_N} > \frac{\gamma K}{r} \left( \mu - \frac{p_N}{p_P} \right):$ $E_{ext}^{MH} > E_{stab}^{PH} \Leftrightarrow \frac{c_N - c_P}{p_N q_N} < K$ $E_{ext}^{MH} > E_{stab}^{NH} \Leftrightarrow \frac{c_N - c_P}{p_N q_N} < \frac{m}{\mu \gamma}$ | $E_{ext}^{NH} > E_{stab}^{NH}$ ⇔ $\frac{c_N - c_P}{p_N q_N} < \frac{m}{\mu \gamma}$ |

It appears that a stable predator-focused system is reached at low efforts (for efforts smaller than 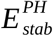), while a stable prey-focused system is reached at high efforts (for efforts larger than 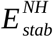). In between, a mixed harvest is reached. Interestingly, the feasibility conditions of the mixed harvest equilibrium are defined by the limit efforts 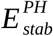 and 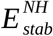 that determine the stability of the predator-focused and the prey-focused systems. If 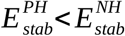, this mixed harvest equilibrium is stable. But if on the contrary 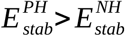, then the equilibrium is unstable. Moreover in this case, three equilibria overlap: stable predator- and prey-focused equilibria, and the unstable mixed harvest equilibrium. The system then exhibits bistability.

In the last row of the table, we investigate the relationships between the feasibility and stability conditions of the different equilibria. Importantly, it appears that if the predator goes extinct in one of the equilibria, then other equilibria cannot be reached for higher efforts. To illustrate this, let us consider three cases, where we suppose that a stable mixed harvest is possible (*q_P_* / *q_N_* >(*γK* /*r*)(*μ* − *p_N_* / *p_P_*)). In the first case, the difference between prey and predator costs relative to prey price and catchability is larger than prey carrying capacity ((*c_N_*−*c_P_*)/ (*p_N_ q_N_*)>*K*). This means that harvesting prey is costly and does not bring much profit relative to its maximum density. Therefore, the interest of harvesting prey is limited. As a result, the predator goes extinct before reaching the mixed equilibrium 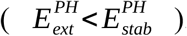. It also implies that the mixed equilibrium is unfeasible 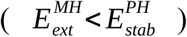. As predator mortality is considered to be lower than its maximum growth rate (*m*<*μγK*) it also implies that the prey-focused equilibrium is unfeasible 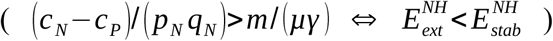.

In the second case, the difference between prey and predator costs relative to prey price and catchability is smaller than prey carrying capacity ((*c_N_*−*c_P_*)/ (*p_N_ q_N_*)< *K*), but larger than predator mortality relative to the effect of predation ((*c_N_*−*c_P_*)/ (*p_N_ q_N_*)>*m*/(*μγ*)). Here the prey gets harvested in the mixed equilibrium, but the predator disappears before reaching the prey-focused equilibrium. Then, the prey-focused equilibrium is unfeasible.

In the third case the difference between prey and predator costs relative to prey price and catchability is smaller than prey carrying capacity ((*c_N_*−*c_P_*)/ (*p_N_ q_N_*)< *K*), and smaller than predator mortality relatively to the effect of predation ((*c_N_*−*c_P_*)/ (*p_N_ q_N_*)<*m*/(*μγ*)). This is always the case when the cost of harvesting prey is smaller than the cost of harvesting predators, which is often true as predators are generally bigger than their prey. In this case, all three equilibria are feasible.

## Appendix B: Robustness of results from the Rosenzweig-MacArthur model

In the main text, we show that increased adaptation speeds can induce oscillations and therefore destabilise the mixed harvest equilibrium of a Rosenzweig-MacArthur system. Here we assess the sensitivity of these results to other parameters of the model. To do so, we numerically compute the Jacobian of our system for each parameter value. When a pair of complex eigenvalues crosses the imaginary axis, the system undergoes a Hopf bifurcation and begins to oscillate.

Figure A2 shows the influence of all parameters of the model on the stability of the mixed harvest equilibrium, relative to adaptation speed. The main conclusion is that increased adaptation speed always displayed a destabilising influence on the mixed harvest equilibrium. Moreover, high adaptation speeds broaden the range of parameters for which oscillations occur. These results therefore support the main text’s conclusion of a destabilising effect of adaptation speed in this model.

The influence of the other parameters can be organised in three categories. Figures A2a, d, g and j show parameters that are destabilising at low values and stabilising at higher values. Interestingly, the total fishing effort shows this kind of pattern, meaning that high fishing efforts can potentially stabilise an oscillating system. Figure A2b, e, h, and k show parameters that destabilise the equilibrium, jointly with adaptation speed. One of these parameters is the carrying capacity, which agrees with the well-known paradox of enrichment [40]. Also note that parameters that favor the relative profitability of prey, namely prey price and predator fishing cost, are destabilising. Figure A2c, f, i, l and m shows stabilising parameters. In accordance with other studies [41], we find that the predator mortality rate is stabilising. Also note that parameters that favor the relative profitability of predators, namely predator price and prey fishing cost, are stabilising.

**Figure A2:**
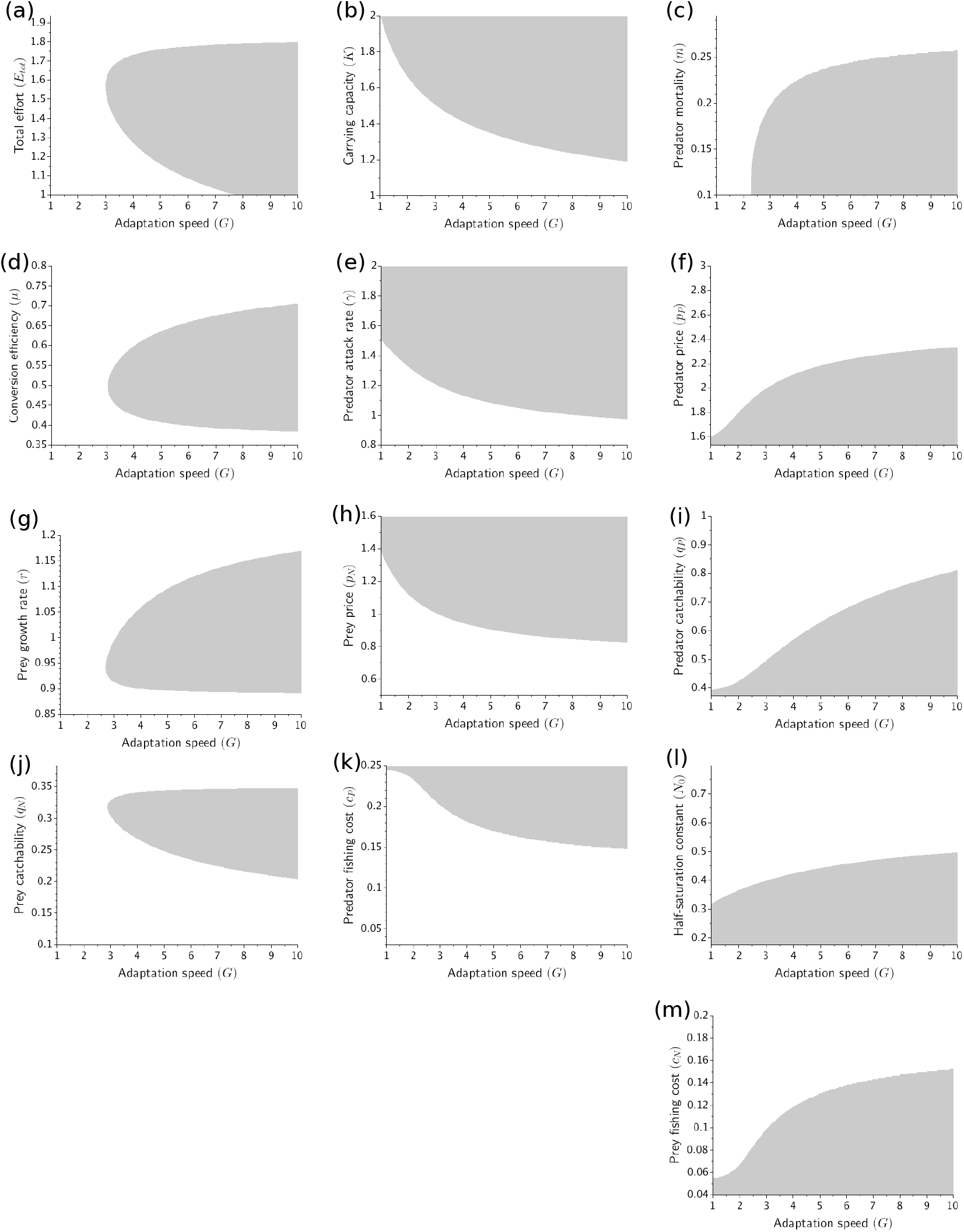
Influence of various parameters on the stability of the mixed harvest equilibrium, relative to adaptation speed. In the white zone, the system is stable, while in the grey zone it oscillates. Value of fixed parameters: *r*=1, *K*=1.5, *m*=0.2, *μ*=0.5, *γ*=1.2, *q*_*N*_=0.3, *q_P_*=0.5, *p*_*N*_=1, *p*_*P*_=2, *c*_*N*_=0.1, *c*_*P*_=0.2, *N*_0_=0.4, *E_tot_*=1.5.

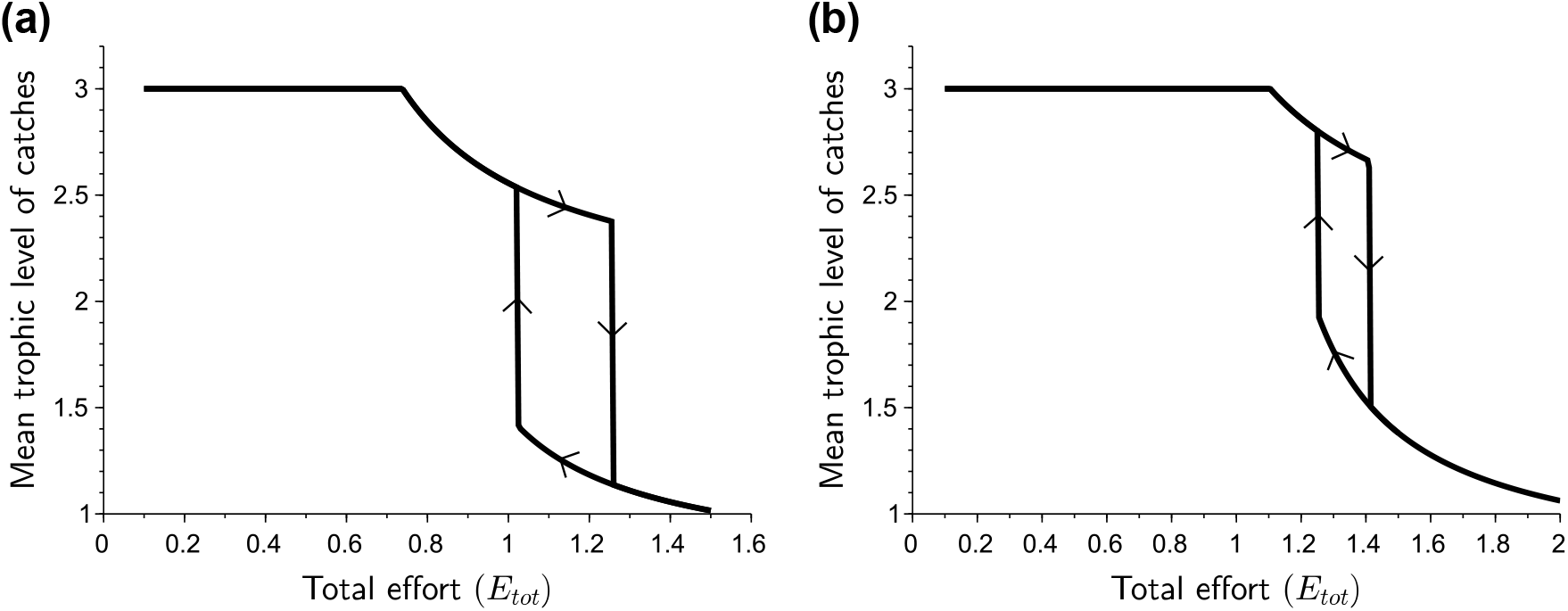

